# A novel machine learning-based approach for the detection and analysis of spontaneous synaptic currents

**DOI:** 10.1101/2021.11.09.467896

**Authors:** Thomas Pircher, Bianca Pircher, Andreas Feigenspan

## Abstract

Spontaneous synaptic activity is a hallmark of neural networks. A thorough description of these synaptic signals is essential for understanding neurotransmitter release and the generation of a postsynaptic response. However, the complexity of synaptic current trajectories has either precluded an in-depth analysis or it has forced human observers to resort to manual or semi-automated approaches based on subjective amplitude and area threshold settings. Both procedures are time-consuming, error-prone and likely affected by human bias. Here, we present three complimentary methods for a fully automated analysis of spontaneous excitatory postsynaptic currents measured in major cell types of the mouse retina and in a primary culture of mouse auditory cortex. Two approaches rely on classical threshold methods, while the third represents a novel machine learning-based algorithm. Comparison with frequently used existing methods demonstrates the suitability of our algorithms for an unbiased and efficient analysis of synaptic signals in the central nervous system.

## Introduction

Until now, the analysis of spontaneous excitatory and inhibitory postsynaptic events in neurons relies to a large extent on the manual inspection and processing of current traces. Although several software packages promise an automated peak detection, the overall process is extremely time consuming, often counterintuitive, and it is mainly based on the expertise of a human observer. Although advanced training skills are combined with high levels of alertness during hours of analysis, consistent results are still difficult to obtain. Especially, the unknown variable of human bias must be taken seriously. People are attributed to have extraordinary abilities with respect to sense-making and reasoning, but they are also susceptible to an innate bias that unconsciously shapes their decision-making [1]. A bias acts like a filter, which funnels information into diverse channels in order to avoid impending sensory overload. However, this filtering can lead to major errors and misinterpretations as it unknowingly infiltrates the analysis of complex signals. For example, a pre-existing hypothesis may influence decision-making and it may make one unconsciously ignore contrary evidence [2]. In psychological science, the so-called anchoring effect is described as the tendency to focus selectively on a subset of the available information when making decisions [3].

In order to eliminate these sources of error in the analysis and subsequent interpretation of spontaneous postsynaptic events, it was our intention to develop algorithms, which make the results reliable and reproducible and simultaneously minimize expenditure of time.

Spontaneous synaptic events in a discrete time series can be described as frequently occurring anomalies. Anomaly detection is a wide area subject to extensive research [4]. A common approach includes ARMA, ARIMA and ARIMAX [5],[6],[7], where the unsupervised learning model will be trained to predict the regular behaviour of the signal in order to determine non-regular events. However, applying these models to time series of spontaneous activity leads to very slow calculations and a rather poor fit to the data, since the models are not able to identify the inherent logic of the signal. Given these drawbacks, it is not possible to use the forecast approach to test for abnormal noise variance. General noise testing methods [8] are also not adequate for the analysis of spontaneous activity time series. The noise in synaptic signals is similar to analytical noise like white or red noise. This neuronal noise results from the binding of neurotransmitter molecules to their cognate receptors, thereby triggering the opening of ion channels and the subsequent diffusion of ions along their electrochemical gradients. Thus, its spectrum and time correlation is complex and cannot be easily simulated. Kernel or rolling value methods also fail to detect events properly in time series of spontaneous activity. Although persistent tests are able to detect the rising edge of an event, calculation costs are high compared to a slope algorithm with preset threshold values. Finally, the change point detection method [9] has a non-intuitive parametrization, and the detected signal change points do not correlate with measured events. Taken together, current methodological approaches summarized in this section do not appear to be adequate for an unbiased automated analysis of spontaneous synaptic events.

Therefore, we generated and compared three different procedures for a computer-based analysis of spontaneous excitatory postsynaptic currents (spEPSCs). Two approaches are based on threshold methods considerably more advanced than those already available, whereas the third approach relies on a machine learning-based algorithm.

## Results

To overcome the potentially error-prone subjective identification of synaptic events, we aimed at developing novel approaches for an automated data analysis. Spontaneous synaptic currents were recorded with the patch-clamp technique from different retinal cell types including horizontal cells, AII amacrine cells, ganglion cells, ON and OFF cone bipolar cells, and rod bipolar cells. Retinal neurons were routinely filled with a fluorescent dye and identified according to their morphology and location in vertical and horizontal slice preparations of the mouse retina. In addition, spontaneous synaptic events were recorded from neurons in a primary culture of mouse auditory cortex after 21 days *in vitro*.

The first method—the so-called “simple” algorithm—is based on a simple threshold approach that detects local extreme values. To identify these as events, their maximum amplitude and preceding slopes are measured and compared to preset threshold values. To ensure consistent results, the recorded signals are processed using a Blackman-Harris kernel with a width of 21 to filter the data by a 50 kHz sampling rate.

The second algorithm is called “slope”, and it is based on the analysis of the positive and negative slopes of an event. We calculate the first derivative of the filtered signal and define positive and negative threshold values. In general, an event is characterized by an inward current, starting with a steep negative slope, which exceeds the preset threshold value and thus activates a negative trigger. Following a local minimum, the positive slope of the decaying phase eventually falls below the threshold, which designates the endpoint. The third, machine learning-based algorithm depends upon unsupervised trained kernels, which describe the state of an event at a certain point in time (“autoclassifier”). Only if the signal passes through all states, it will be declared as an event. Figure 1a displays these states as windows of 128 time steps assigned to different clusters. The labels of the clusters are randomly chosen by the k-means algorithm, whereas the order corresponds to the event age. It should be noted that this figure merely approximates a visual representation, as the information of the latent domain can only be interpreted by the classifier but not by a human observer. The autoclassifier is based on features, not on shapes. Nevertheless, internal processes of this method can be depicted quite clearly in this way, thus providing some insight into how the black box of the autoclassifier works. Figure 1b shows the dimension-reduced autoencoder information with colours assigned to the clusters outlined in Figure 1a. We reduced the latent domain of the autoencoder with the unsupervised umap algorithm [10], which only affects the latent domain information but not the assigned clusters, from originally eight to two dimensions. Interestingly, the resulting projection shows a regular organization: The radius of a polar coordinate system centered on the projection functions as a separation criterion for noise, whereas its angle corresponds to the age of the event in the analysed time window (Fig. 1b). The border between the first and last cluster is located at *θ* ≈ 265, and the event state ages clockwise from cluster 4 to 1. Since the reduction from eight to two dimensions involves a loss of information, clusters 3 and 4 appear blurred, and cluster 5 partly contains areas from other clusters. We refer to the Methods section for a more detailed description of signal processing and classification algorithms.

**Fig 1.**
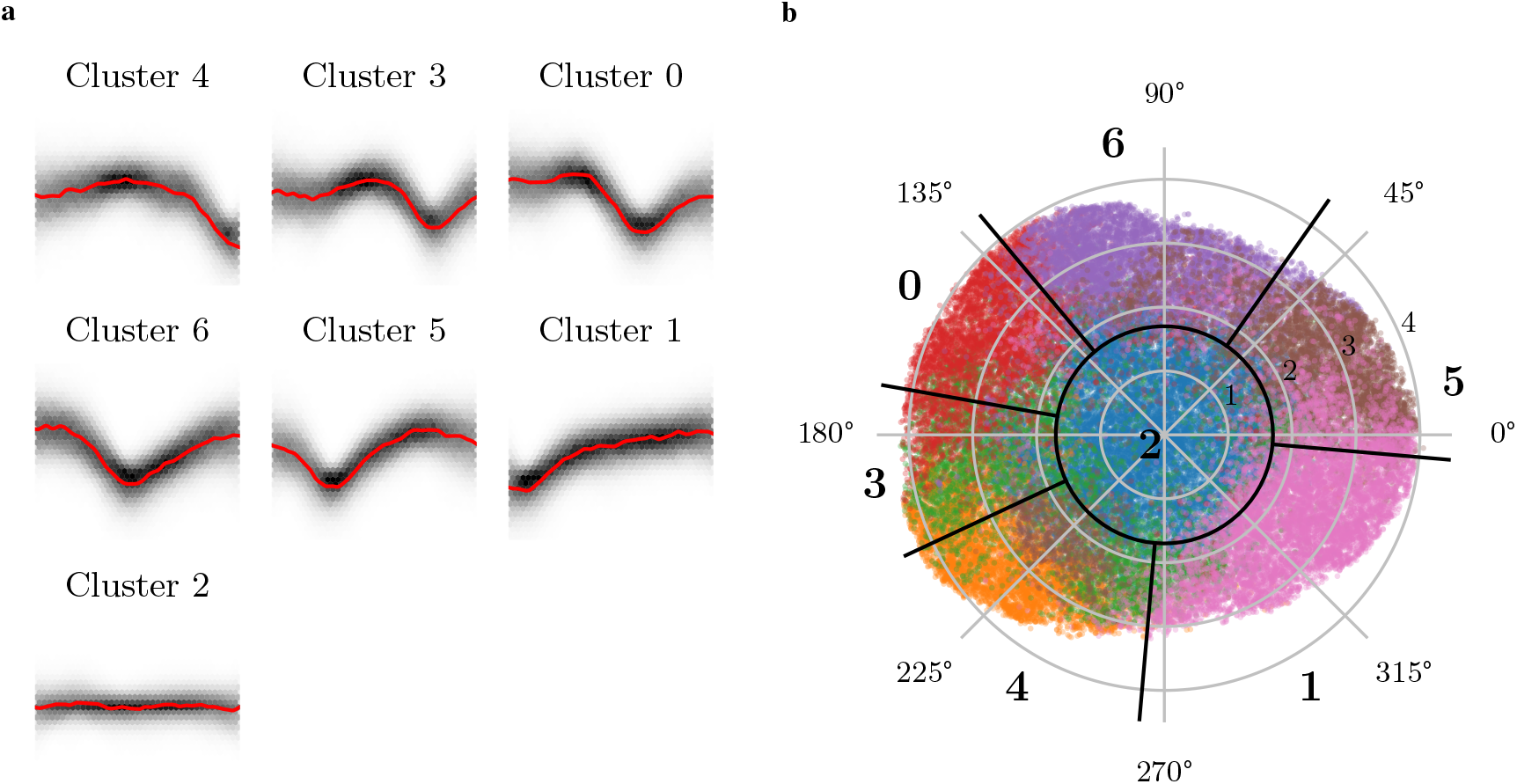
a Signal density and common signal of the different clusters. All graphs have the same scale. The signals comprise 128 time steps, each time step being 10 *µ*s. The graphs are sorted in chronological order. The shaded areas show the density of data points, and the red line represents the approximated shape of the state based on a Nadaraya-Watson-estimator. **b** Dimension-reduced latent domain generated by the umap algorithm. The 2D umap values are mapped to polar coordinates with a center at *x*_1_ = 4.95 and *x*_2_ = 4.1. The colours represent the different clusters that were determined with the k-means algorithm. Radial black lines indicate the approximated borders of the clusters. The blue cluster in the centre (**2**) represents windows containing only noise. Starting with cluster **4**, the chronological order of the clusters in **a** is represented as a clockwise rotation.

To evaluate the performance and compare the results of these newly developed algorithms, we analyzed the same synaptic current trajectories with four other methods. First, we used the commercial software MiniAnalysis (Synaptosoft Inc., Fort Lee, NJ) in a manual detection and a semi-automated analysis mode, which we will refer to as “ma-manual” and “ma-auto”, respectively. Second, we used the multi-step threshold algorithm of the software Neuromatic [11], based on the work of Kudoh and Taguchi [12] (designated as “kudoh”). Finally, we employed the threshold-based algorithm by Muthmann et al. [13] that was originally designed to detect events in multi-electrode array (MEA) recordings (designated as “muthmann”).

Figure 2 shows a representative current trace from a retinal horizontal cell (a) and a cell from cortex primary culture (b), each displaying multiple synaptic events. The positions of the events detected by each algorithm are indicated by vertical lines. Both signals clearly differ with regard to amplitude and event duration, making it rather difficult to implement a generally applicable algorithm that is not trained on a specific cell type. Large events were detected by all methods, although some jitter with respect to timing was apparent in each case. In contrast, smaller events lacking a well-defined baseline were identified only by some methods. Time series of horizontal cells usually displayed a rather irregular sequence of synaptic events with changing amplitudes and kinetics, and a clearly discernible baseline was mostly lacking. As a consequence, it was especially difficult in this cell type to determine the beginning of an event, to measure its peak amplitude with respect to a variable baseline, and to reliably distinguish between signals and noise. Thus, it is not surprising that the different methods show a slightly larger deviation in the number and positions of events found in signals from horizontal cells as compared to those of cortical neurons. However, the three newly developed algorithms show a relatively high concordance for both cell types. In contrast, the “muthmann” algorithm originally used for MEAs clearly reveals its limitations when applied to horizontal cells. In most of the datasets, this approach misses distinct events resulting in a reduced number of peaks. The output of the “kudoh” algorithm is also considerably different. This algorithm interprets virtually every current fluctuation in horizontal cells as an event, whereas it fails to detect obvious events in cortical cells. In these cells, the “simple” algorithm also detects relatively few events. This is caused by the larger range of event amplitudes, and it could be improved by optimized filtering before detecting local extreme values.

**Fig 2.**
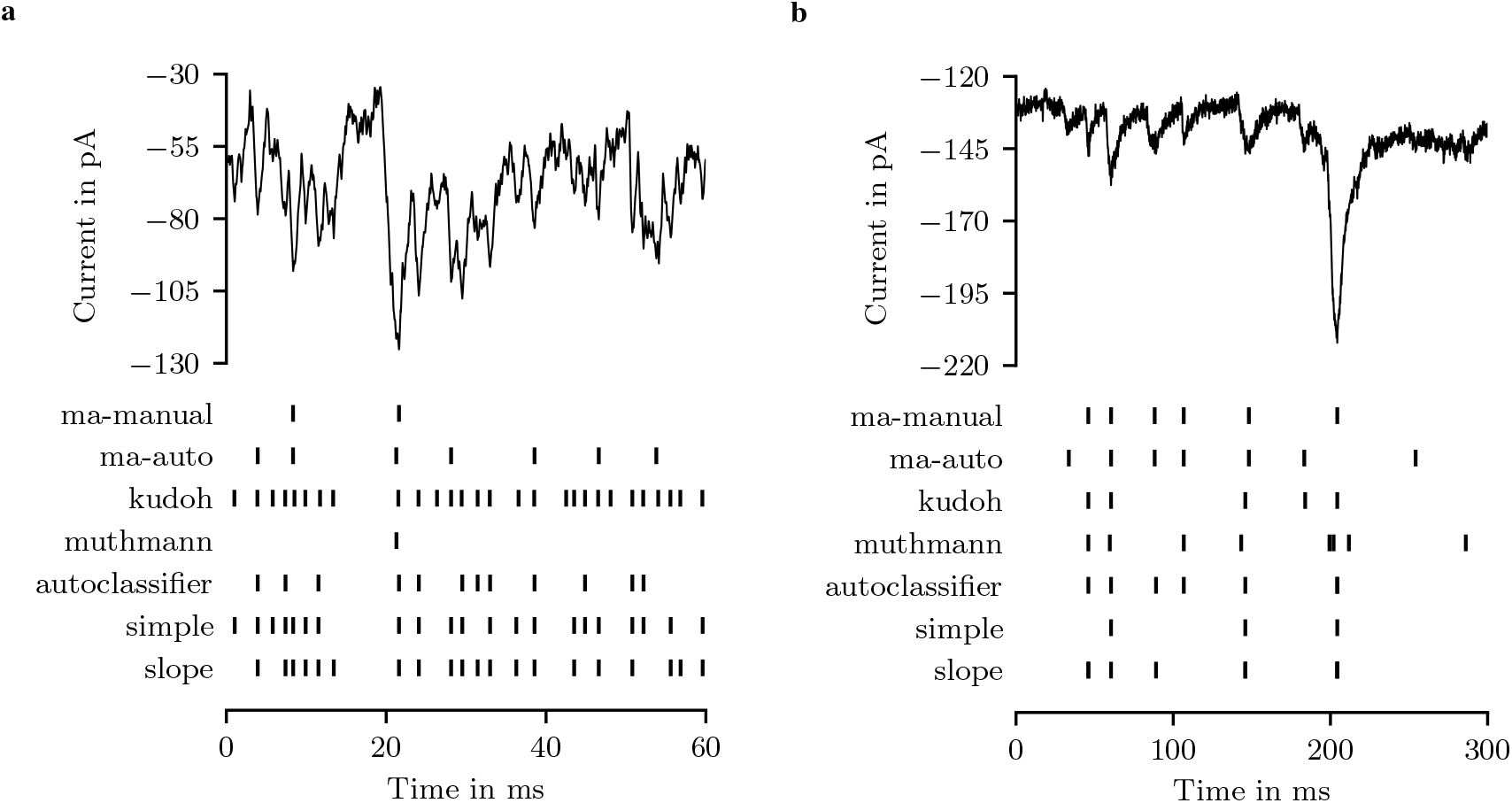
Representative time series of spontaneous synaptic activity measured in (**a**) a retinal horizontal cell and (**b**) a cultured cortical neuron (upper panel), and peak positions of events detected by the algorithms specified on the left (lower panel). “ma-manual” and “ma-auto” were carried out with the MiniAnalysis software package, “kudoh” refers to the multi-step threshold algorithm of [12], and “muthmann” to the MEA software package ([13]). “autoclassifer”, “simple” and “slope” are novel algorithms described in the main text.

Figure 3 provides an overview of event detection performances of all algorithms used in this study. The matrices represent the percentages of missed events in direct comparison between the algorithms for all cell types. Darker shades represent a low coverage, whereas lighter shades indicate highly concordant analysis results. It is rather complicated for any algorithm to fully cover the event characteristics of these diverse cell types. Additionally, the lack of a ground truth for cell types with a low signal-to-noise ratio makes the validation of the accuracy of the algorithms difficult. False positives increase the missing rates of the remaining algorithms. In most cases, the algorithms “slope”, “simple” and “autoclassifier” show quite similar results, although the “autoclassifier” generally detects fewer events. As compared to the other algorithms, “threshold” and “ma-auto” differ most, suggesting a rather high error rate in event detection. Apparently, “ma-auto” often does not recognise obvious events. In some cases, the suggested peak position is neither at the minimum, nor do the other parameter values correspond to the expected shape of an event. Even through numerous iterations in the manual adjustment of parameters, ideal values could not be determined. Of course, continuous readjustment makes the process of manual evaluation more difficult and time-consuming.

**Fig 3.**
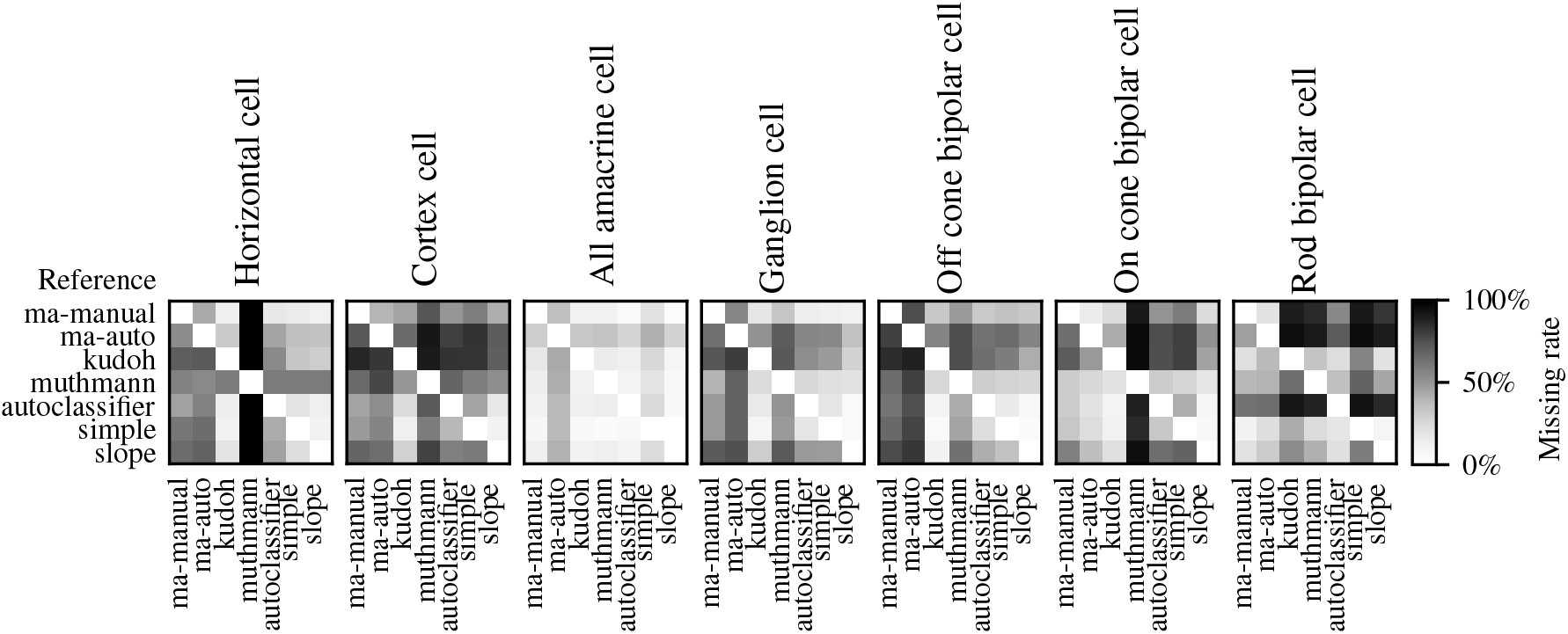
Comparison of algorithm performances in event detection. The matrices show the rates of events missed by an algorithm related to a reference for various cell types.”ma-manual” and “ma-auto” were carried out with the MiniAnalysis software package, “kudoh” refers to the multi-step threshold algorithm of [12], and “muthmann” to the MEA software package ([13]). “autoclassifer”, “simple” and “slope” are novel algorithms described in the main text.

Table 1 specifies the mean number of events per current trace detected by each algorithm and its speed relative to manual detection for retinal horizontal cells and cultured cortical neurons. The number of events is calculated as the mean of all traces under study. As the length of individual current traces as well as the mean duration of events differ between horizontal cells and cortical neurons, the count of mean events is different between the two datasets. To evaluate the performance of each algorithm with respect to manual analysis, its speed is measured and normalized to the time necessary for manual detection (estimated as 10 min per 5 s current trace). In horizontal cells, the “slope” algorithm identifies most events, followed by “simple” and “autoclassifier”. These algorithms detect near to or more than twice as many events compared to the “ma-manual” approach. The threshold-based algorithms “kudoh” and “muthmann” perform rather poorly in the analysis of complex horizontal cell current trajectories with 1333 and 1309 events per second, respectively (see also Fig. 3). When a stable baseline is present as in recordings of cultured cortical neurons, the divergence of algorithm performances appears not as pronounced. With respect to “ma-manual”, the “slope” algorithm displays the fastest performance, whereas the “autoclassifier” is relatively slow. The “autoclassifier” calculates the state for every time step, which is a convolution-like operation and therefore computationally expensive. Thus, the “autoclassifier” is always slower than both non-machine learning algorithms, but it is still considerably faster than “ma-manual” detection.

**Table 1.** Table showing the mean number of events and the relative speed of the specified algorithms normalized to “ma-manual” analysis.

|  | horizontal cell |  |  | cortical neuron |  |  |
| --- | --- | --- | --- | --- | --- | --- |
|  | mean<br>events / s | relative speed<br>to length | to events | mean<br>events / s | relative speed<br>to length | to events |
| **ma-manual** | 474.1 | 1 | 1 | 13.84 | 1 | 1 |
| **ma-auto** | 576.9 | 1 / 214 | 1 / 260 | 28.98 | 1 / 36 | 1 / 76 |
| **kudoh** | 1333 | 1 / 50 | 1 / 139 | 60.17 | 1 / 10 | 1 / 43 |
| **muthmann** | 1309 | 1 / 74 | > 1 | 8.244 | 1 / 169 | 1 / 101 |
| **autoclassifier** | 759.2 | 1 / 2 | 1 / 3 | 11.97 | 1 / 4 | 1 / 4 |
| **simple** | 984.5 | 1 / 375 | 1 / 779 | 10.85 | 1 / 921 | 1 / 722 |
| **slope** | 1208 | 1 / 15 | 1 / 38 | 21.49 | 1 / 974 | 1 / 1513 |

For a more comprehensive description of detected events, we measured duration, peak amplitude, 10% −90 % rise time, and time constant of exponential decay of individual spEPSCs. Figure 4 shows the respective kernel density distributions for events identified with the algoritms “slope”, “autoclassifier”, and “ma-manual”. All other algorithms do not provide these parameters, and therefore they are not considered here. We decided to display the data as kernel density distributions instead of histograms, since the choice of bin width influences the shape and thus the interpretation of the distributions.

**Fig 4.**
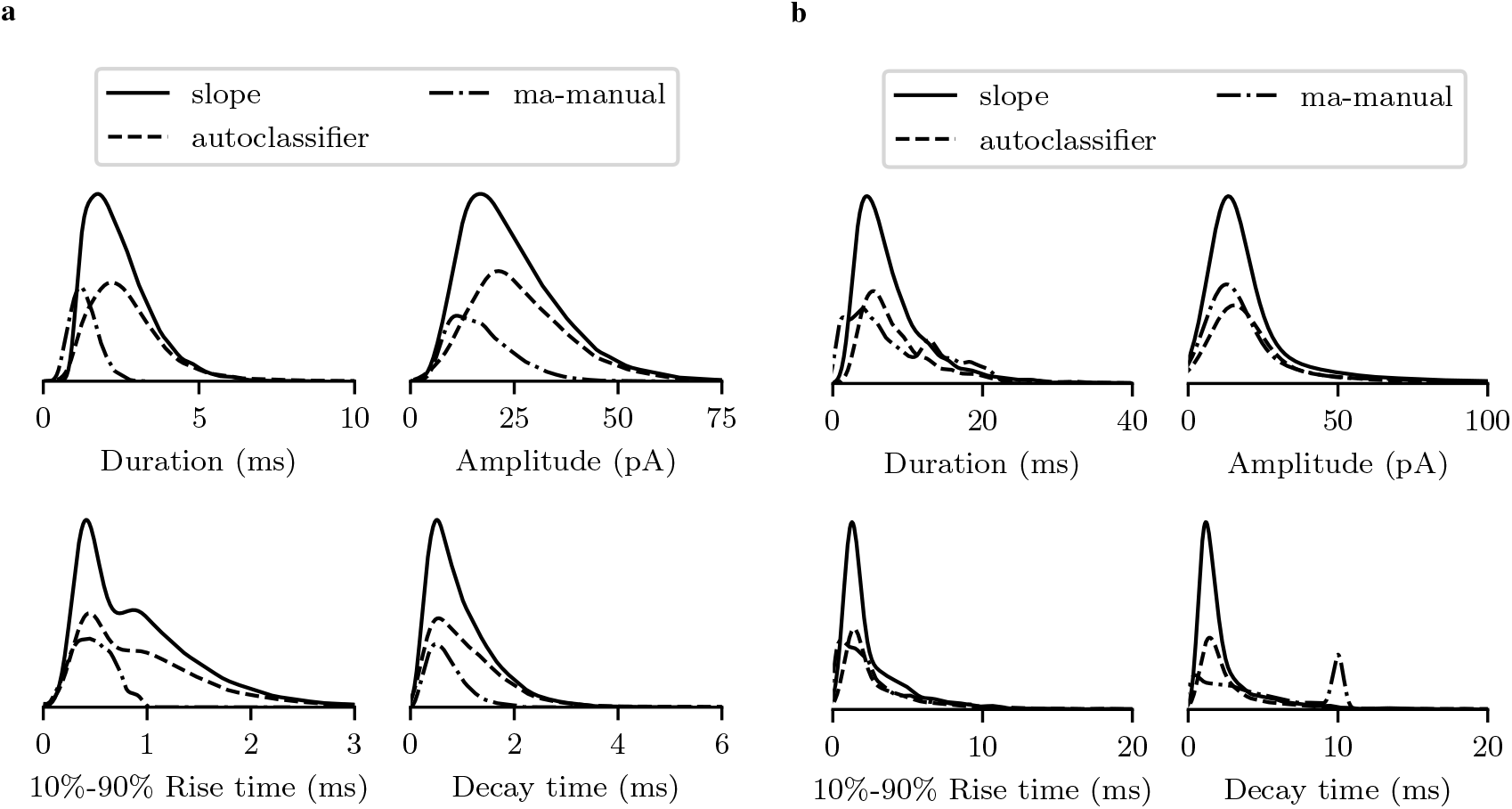
Kernel density distributions of the event parameters duration, amplitude, 10%-90% rise time and time constant of decay as determined by “ma-manual”, “autoclassifier”, and “slope” algorithms for horizontal cells (**a**) and cultured cortical neurons (**b**). Since the sum of the integrals of the distributions is scaled to 1, the areas represent the set size of each distribution related to the others.

In case of horizontal cells, we observe major differences between manual and automated approaches (Fig. 4a). Manually detected events are characterized by small amplitudes and rather fast kinetics, whereas events identified by the “autoclassifier” algorithm appear larger with slower kinetics (see the pronounced tail in the 10% −90% rise time and decay time distributions). With respect to duration and amplitude, results obtained with the “slope” algorithm are located between the two other approaches, whereas they are more similar to the “autoclassifier” concerning rise and decay times.

Differences between manual and automated detection appear less pronounced for cultured cortical neurons, which display a rather distinct baseline. The second peak at around 10 ms in the decay time distribution is likely attributable to a computing error in the software for the manual detection. To obtain a valid peak shape, parameters must be adjusted in a way that the decay time occasionally overlaps with the following rise time. Therefore, this second peak is considered an artefact.

## Discussion

In this study, we present three novel algorithms for a fully automated analysis of spontaneous excitatory postsynaptic currents measured in several cell types of the mouse retina as well as in a primary culture of mouse auditory cortex. Two approaches rely on considerably advanced threshold methods, whereas the third approach is based on machine learning algorithms.

In principle, it is rather difficult to evaluate algorithms, which apparently have no predetermined “correct” solution. By analysing spontaneous synaptic activity, it is impossible to reliably distinguish between noise and a true neuronal response caused by release of a single presynaptic vesicle. Detected events might include false positives, whereas actual events could also mistakenly be defined as noise. Nevertheless, the algorithms presented here outperform already existing and previously used methods. In addition to a fast and unbiased approach, which most of the other algorithms also accomplish, our event detection is very robust. Events are reliably and consistently identified and analysed across the entire data set comprising several cell types from distinct regions of the central nervous system. Other algorithms, especially “muthmann”, which was the most unreliable in our tests, repeatedly produced time windows with no events despite visual evidence to the contrary. Additionally, third-party algorithms are often quite sensitive to noise, and they utilise a counterintuitive and/or time consuming trial and error-based procedure to determine appropriate detection parameters. This concerns in particular the software package Mini Analysis, which produced the highest rate of false positives in the semi-automated analysis.

Algorithms already in common use are compared with the three novel approached described in this study (see Table 2). Thus, for example, the “autoclassifier” algorithm is completely insensitive to noise, which eliminates the need for potentially error-prone filtering. The algorithm is highly reliable, and its inherent logic based on feature-driven detection distinguishes this approach from the other algorithms. Since the “autoclassifier” is trained unsupervised, its decision making is completely unbiased. Only special hardware and software requirements as well as the necessary expertise for training slightly limits the general application of this machine learning algorithm. Simple and very robust in its application and also particularly intuitive is the parametrization of both “simple” and “slope” algorithms. However, the most critical parameter here concerns the lower limit of the signal amplitude, above which a deviation from baseline will be considered an event. This parameter, which likely corresponds to the release of a single vesicle from a single active zone, could not be reliably estimated with any of the algorithms investigated.

**Table 2.**
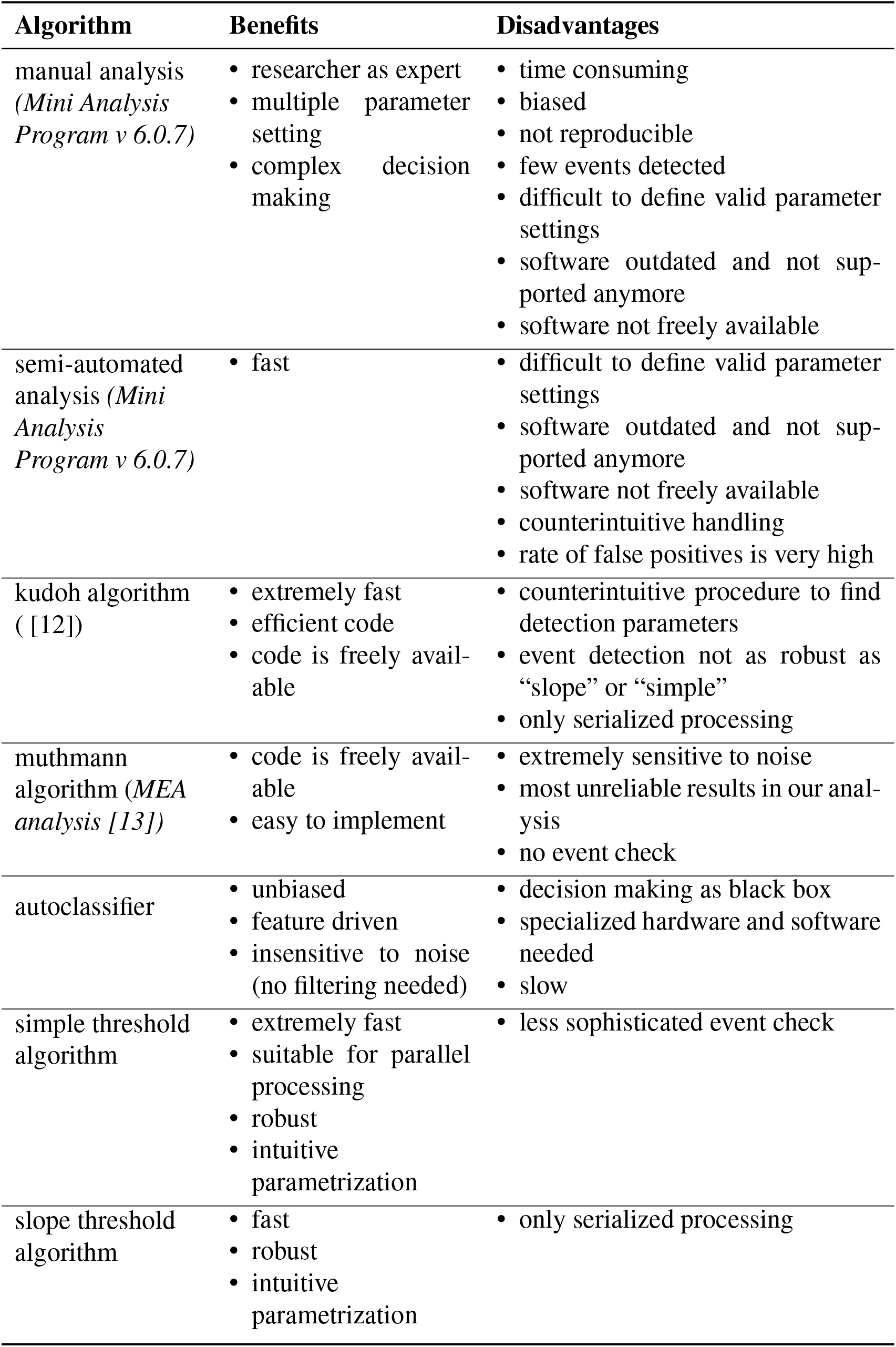
Overview of algorithms applied for analysis of spEPSCs.

| Algorithm | Benefits | Disadvantages |
| --- | --- | --- |
| manual analysis<br>(*Mini Analysis Program v 6.0.7*) | • researcher as expert<br>• multiple parameter setting<br>• complex decision making | • time consuming<br>• biased<br>• not reproducible<br>• few events detected<br>• difficult to define valid parameter settings<br>• software outdated and not supported anymore<br>• software not freely available |
| semi-automated analysis (*Mini Analysis Program v 6.0.7*) | • fast | • difficult to define valid parameter settings<br>• software outdated and not supported anymore<br>• software not freely available<br>• counterintuitive handling<br>• rate of false positives is very high |
| kudoh algorithm<br>( [12]) | • extremely fast<br>• efficient code<br>• code is freely available | • counterintuitive procedure to find detection parameters<br>• event detection not as robust as “slope” or “simple”<br>• only serialized processing |
| muthmann algorithm (*MEA analysis [13]*) | • code is freely available<br>• easy to implement | • extremely sensitive to noise<br>• most unreliable results in our analysis<br>• no event check |
| autoclassifier | • unbiased<br>• feature driven<br>• insensitive to noise (no filtering needed) | • decision making as black box<br>• specialized hardware and software needed<br>• slow |
| simple threshold algorithm | • extremely fast<br>• suitable for parallel processing<br>• robust<br>• intuitive parametrization | • less sophisticated event check |
| slope threshold algorithm | • fast<br>• robust<br>• intuitive parametrization | • only serialized processing |

Across the cell types under investigation, we find a considerably higher agreement between our novel algorithms than between the manual and the computer-based methods. This, of course, is most likely due to the previously described bias, which inevitably occurs during manual evaluation. This human bias impedes the reproducibility of the results carried out by different researchers, and it therefore clearly reduces the reliability and significance of the analysis.

With the exception of horizontal cells, which only receive excitatory glutamatergic input from photoreceptors, all other neuronal cell types investigated here receive both excitatory and inhibitory synaptic input. However, the algorithms presented in this study only consider excitatory, but not inhibitory currents. Inhibitory events are significantly longer [14], exceeding the 128 time steps of the “autoclassifier”, and thus they need to be analysed with a different parametrization. Similarly, the “slope” algorithm has a preset value for the maximum duration of an event. The possibility to also analyse inhibitory currents, and multi-channel recordings with extended methodologies, will be addressed in prospective versions of the algorithms. Finally, we would like to introduce the option of an easy-access application of the algorithms in order to make them applicable for the analysis of synaptic signals independent of computational expertise.

## Methods

In this study, we compare novel event-detecting algorithms with already existing methods. Therefore, we briefly describe the workings of the Mini Analysis Program (“ma-manual” and “ma-auto”), the Neuromatics software (“kudoh”), and a software package dedicated to the analysis of MEA signals (“muthmann”). Our newly developed algorithms are described in greater detail thereafter.

### Manual and semi-automated detection with the software package *Mini Analysis Program Version 6*.*0*.*7*

Detection parameters were adjusted manually following preset values including amplitude and area threshold, the period to search for a local maximum and decay time, and the stretch of baseline before a peak. The parameters were chosen and optimised to match visual characteristics of spontaneous currents. We distinguished between an entirely manual analysis, in which each peak was identified and selected individually by eye, and a semi-automatic analysis. The latter was carried out using the “Non-Stop Analysis” function with suitable parameters settings. The same set of parameters was used for both manual and semi-automatic analysis. In addition, a manual correction of the semi-automatic results was omitted in order to ensure the comparability of the results.

### Kudoh algorithm

Since selling and support of the MiniAnalysis software package has been aborted, the software Neuromatic [11] is now widely used. The event-detecting algorithm is based on a multi-step threshold mechanism [12]. The algorithm iterates over every time step and waits for a forward slope trigger. If the initial search indicates an event, the on set point is set by iterating over the time steps until the signal value exceeds a predetermined noise level. Starting from the on set point, the signal peak is determined by a forward iteration followed by a backward iteration. The peak detection compares mean values in the near vicinity to a reference point, leading to an approximation of the peak position. With increasing noise, however, the peak detection deteriorates significantly. Finally, kinetic parameter are calculated by fitting of a multi-exponential function based on [15]. We did not carry out this fitting procedure, since we were only interested in event detection.

### Muthmann algorithm

For analysing data obtained with micro electrode arrays (MEAs), a threshold-based detection approach was developed. This algorithm searches the baseline of the signal and approximates the noise level statistically [13][16]. An event will be detected if the signal exceeds the predetermined noise level. We used a threshold factor of 4 with respect to the median absolute deviation of the band pass-filtered signal. Current traces were preprocessed with a 0.1–2 kHz 3-pole Bessel filter before analysing the baseline and the noise level (see [13]).

### Autoclassifier

In general, an autoencoder produces an unsupervised transformation of a dataset to an abstract image domain. We extended this concept to allow for a state-based interpretation of synaptic signals, which is completely free of bias and only depends on its own characteristics. Fig. 5 shows the concept of our autoencoder extension, which we termed autoclassifier.

**Fig 5.**
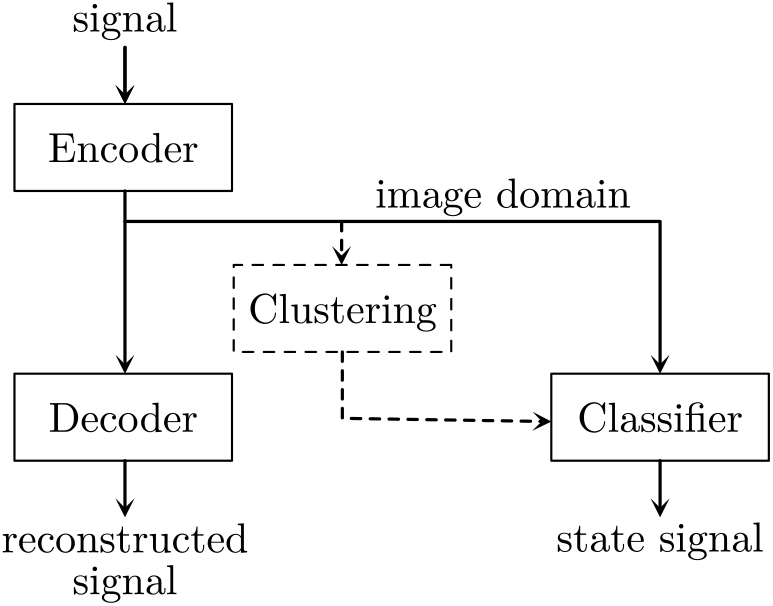
Schematic illustration of the autoclassifier concept.

The image domain of an autoencoder represents the trained states of a signal, whereas the decoder reconstructs the signal from this domain. Thus, it is mandatory that the image domain contains all necessary information. Encoder and decoder have to agree in the interpretation of the signal. The resulting domain necessarily represents the signal information as an abstract logic. A clustering of sample data in the image domain results in groups of common states, and a classifier will be trained to identify these states. The combination of encoder and classifier interprets short sections of the current signal as states, which are further analysed.

The autoencoder and classifier are trained with supervised algorithms, but the autoclassifier creates its inherent logic unsupervised, like an autoencoder.

#### Autoencoder

Encoder and decoder are constructed with three feedforward layers. A window of 128 data points with a sample rate of 25 kHz was used, so the input of the autoencoder represents 5.12 ms of the signal. Brief events fit completely in this window, whereas longer events extend beyond this time frame, but can be analysed as well. This is limited to an event duration of maximal 3 times as long as an average trained event.

Figure 6a shows the structure of the autoencoder. All layers are dense layers; thus, the layers are fully connected. The encoder starts with 1024 followed by 256 neurons.

**Fig 6.**
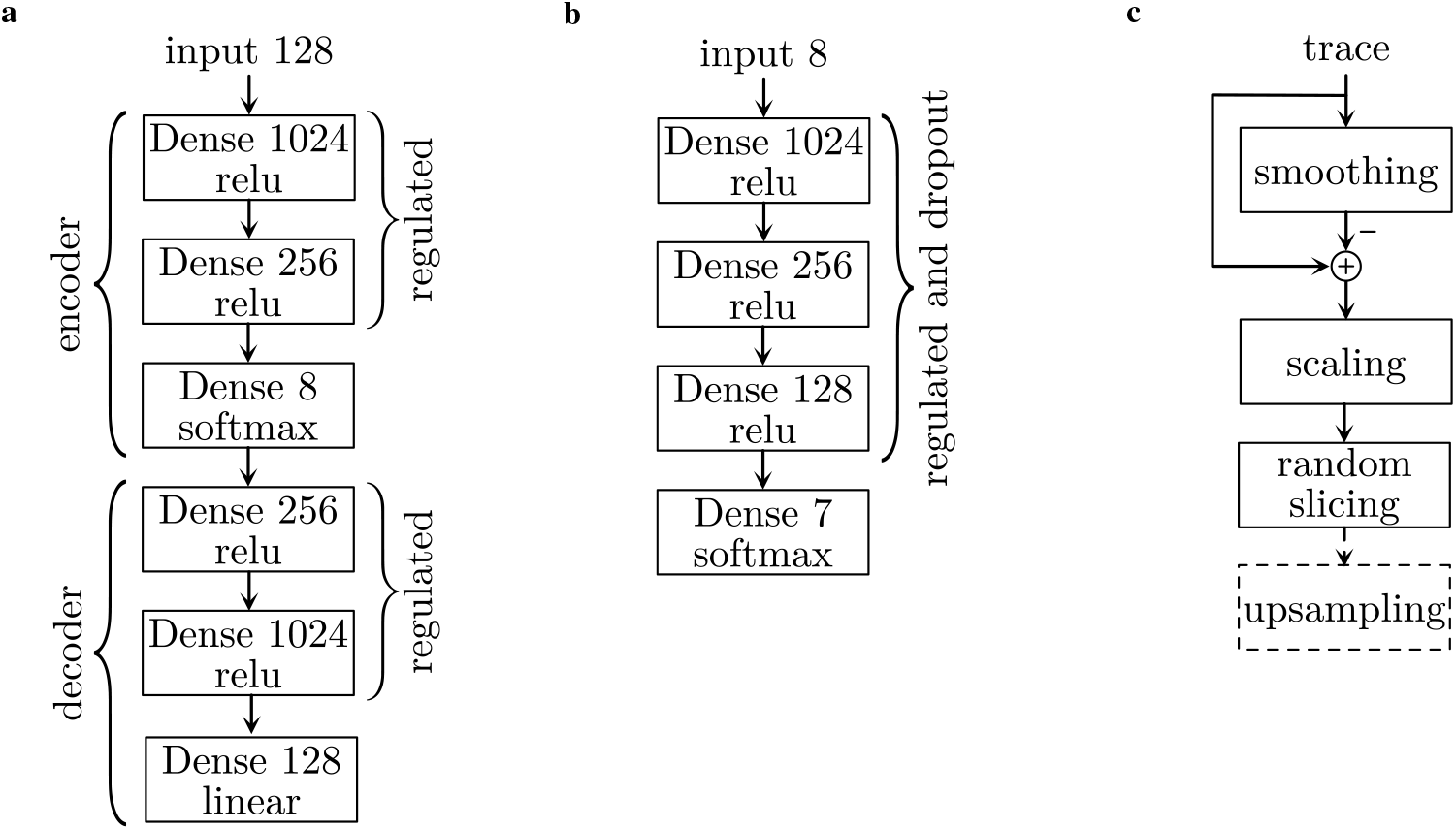
Schematics of the components of an autoclassifier and the data preprocess. **a** autoencoder model, **b** classifier model, **c** data preprocess

Both layers use the “relu” activation. To provoke a sparse layout of the connections, both layers are regulated with the L1 norm with a value of 5·10^−6^. The final layer of the encoder consists of 8 neurons which are “softmax”-activated and not regulated. The decoder mirrors the structure of the encoder. The first two layers are “relu”-activated and regulated as the encoder layer. The last layer has 128 neurons that are linearly activated to allow the reconstruction of the signal window.

The autoencoder is the combination of encoder and decoder. The dataset taken from [17] was preprocessed and randomly cut into 50·10^3^ windows with a length of 128 time steps each. The encoder was trained for 100 epochs with “ADAM” as optimizer with an initial learning rate of 1 ·10^−3^, “Huber” function as loss and a batch size of 32· 10 ·10^3^ addition random sample windows were used for validation. The learning process was tracked after each epoch. If the model did non improve within 10 epochs, the learning rate was reduced by the factor 0.2 until a minimum learning rate of 1 · 10^−6^ was reached. The learning process resulted in a loss of 3.3877 · 10^−4^ and a validation loss of 2.5864 · 10^−4^.

### Clustering

The autoencoder transforms a signal window of 128 time steps into an eight-dimensional domain and then it inverts this transformation to reconstruct the original signal. To reconstruct the signal based on compressed information, the image domain needs to be systematic. We assume that there is a finite number of states that can be identified by clustering.

From the training data of the autoencoder, 60 · 10^3^ sample windows were randomly chosen and transformed to the image domain by the encoder. The resulting samples were scaled by a robust scaler that removed the median and scaled the data according to the interquartile range. These scaled samples build the database for clustering.

It is possible to interpret the clustering directly based on the autoencoder. Grouped by the labels, the clusters can be interpreted as time series. A Nadaraya-Watson-Estimator creates a hypothetical common time series for a group. The mean of the L2 norm between the common time series and the clustered ones is a value of the fit. The cluster algorithm and the number of clusters was not chosen by an interpretation of the position similarity in the image domain, but rather by the similarity of the time series. The combination of the k-means algorithm with seven clusters resulted in an adequate fit with a small number of clusters. By increasing the number of clusters, the resulting difference between the samples and the common time series will decrease until the number of clusters reaches the number of samples, and the difference disappears.

Figure 1a shows the resulting clusters. The common signal was estimated using the Nadaraya-Watson algorithm with a kernel width of 2.0 related to 128 time steps. Table 3 summarizes the number of samples and the resulting root mean square values (RMS), which are based on the preprocessed data for the machine learning, of the difference between the samples and the common signal. The clusters can be ordered in a way that they show an event in different time steps. In this specific case, the image domain of the autoencoder can be interpreted as the time-related position of the peak in the analysed window. The clusters 4, 3, 0, 6 and 5 are equally distributed and show similar RMS values. Cluster 1 occurs more often and has a smaller RMS value. Based on the shape of the common signal of cluster 1, this cluster could also be interpreted as a rising noise signal. Cluster 2 represents the time window without any event, which is the most frequent case. The lack of a specific signal results in a reduced RMS value.

**Table 3.** Data of the different clusters.

| order | cluster | count | RMS |
| --- | --- | --- | --- |
| noise | 2 | 17275 | 0.048 |
| 1 | 4 | 6538 | 0.073 |
| 2 | 3 | 6699 | 0.073 |
| 3 | 0 | 6490 | 0.075 |
| 4 | 6 | 6930 | 0.073 |
| 5 | 5 | 6912 | 0.071 |
| 6 | 1 | 9156 | 0.066 |

### Classifier

The classifier is build as a simple feed-forward model with three hidden layers, 8 input and 7 output variables (Fig. 6b). The hidden layers use “relu” as activation, they are regulated with the L1 norm and a value of 1 ·10^−3^, and 30% of the weights were random dropout. This prevents memorization in the overparameterized model and should provoke parallel and sparse solution paths in the network. The output layer use “softmax” as activation. The model is optimized with the “ADAM”-algorithm with an initial learning rate of 1 · 10^−3^ for 100 epochs with a batch size of 32. The categorical crossentropy was used as loss value.If the optimizer reaches a plateau, the learning rate was reduced by the factor 0.2 until a minimum learning rate of 10^−6^ was reached. The 60 ·10^3^ data samples of the clustering, which were extended from the label information, were randomly split into 80% training and 20% validation data. These results in a training accuracy of 93.69% and a validation accuracy of 98.00%.

The trained classifier interprets the image domain of the encoder and returns the corresponding label related to the clustering. The encoder and the classifier can be combined like the encoder with the decoder. The resulting autoclassifier interprets a time series window as one of the labels that are founded by the clustering.

### Data acquisition

We performed patch-clamp recordings of spontaneous synaptic events from different retinal cell types including horizontal cells, AII amacrine cells, ganglion cells, ON and OFF cone bipolar cells, and rod bipolar cells as described previously [18]. For identification, retinal cell types were routinely filled with a fluorescent dye and characterized according to their morphology and location in vertical and horizontal slice preparations of the mouse retina. In addition, spontaneous synaptic events were recorded from neurons in a primary culture of mouse auditory cortex after 21 days *in vitro*.

Each data set had to be split and preprocessed before synaptic events could be analysed. To prevent interference of the data, we split the data sets based on the measured sequences. Each cell was measured with a fix automated sequence of multiple signal recording. These sequences were randomly separated in to 80% training and 20% validating. Based on this, every test data is like a new measurement and is independent of the training data. In some cases, the current trace displayed a floating baseline that needed to be adjusted. Current traces were smoothed by a kernel with ten times the length of the time window to be analysed. By subtracting the smoothed time series, the baseline was corrected. Assuming that all traces were sufficiently long to include at least one large event, the trace was scaled to a range of [0 … 1]. Then, the traces were divided into sections of a window size of 128 time steps. To reduce the amount of data to be processed to a useful level, sections were chosen randomly: 50 · 10^3^ random slices of the training cells for training and 10·10^3^ random slices of the validation cells for testing. Figure 6c shows the schematic of data handling. The autoencoder was trained to use a series of 128 time steps with a sampling rate of 25 kHz. In case of a different sampling rate, the slices were resampled accordingly.

### Event detection

For event detection, the systematic signal generated by the autoclassifier had to be interpreted. By sorting the clusters, an event should generate an ascending stair with a final step at the end (Fig. 7a, reverse state). The end of an event exists if a step can be detected. The start of an event is determined by reversing the cluster order. The state signal starts at an event with a step followed by a descending stair (Fig. 7a, forward state).

**Fig 7.**
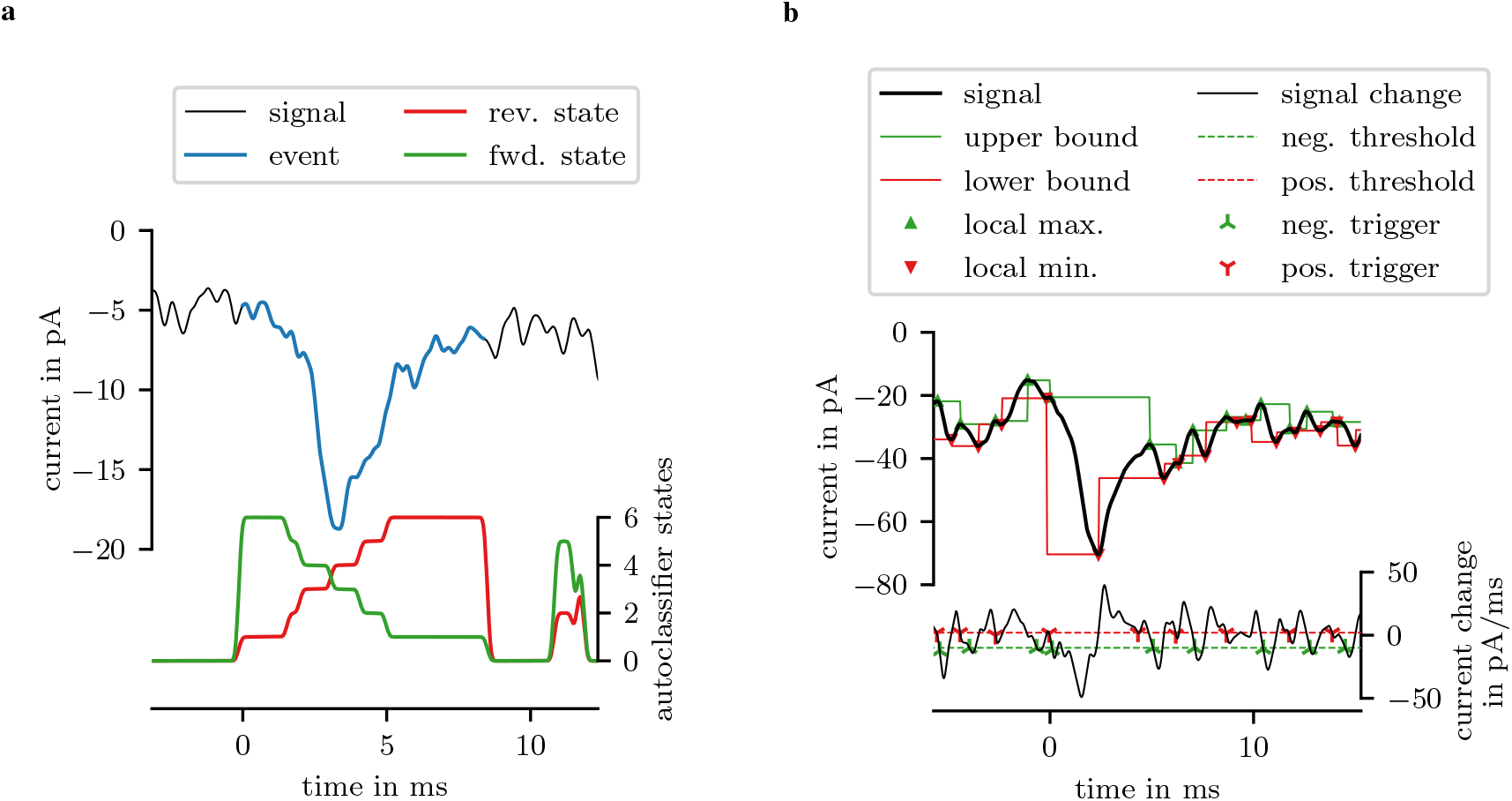
Examples of event detection with autoclassifier and threshold based method.

An event exists in the signal between a starting and an ending points, triggered by the forward and reverse state signal. To compensate the noise in the state signals, these signals were filtered with a short Blackman-Harris kernel of the length 17 time steps. The step was detected, if a slope of 0.51/step (based on the preprocessed data) was reached. In some cases, a start point but no end point was triggered. To exclude these cases, events with a duration greater than 10 ms were removed. Also, events with a duration shorter than 0.8 ms were removed. Additionally, an event was considered valid if the proportion of the noise cluster was less than 25%.

Two non-machine learning based methods were used as a reference to validate the autoclassifier-based approach. The first method detected events by local extreme values as shown in figure 7b. The step of the event had to exist between a local maximum and a local minimum. The signal had to be filtered, since the local extreme values relayed on the noise of the measured signal. An event step will be detected if the peak amplitude and the maximal slope of the step exceed a threshold. This method can not detect the end of an event.

The second non machine learning method based on the direct detection of the slopes. An event starts with a step. These steep slope are characteristic. By reaching the slope trigger, the step of an event will be detected. After the step follows a recuperation in form of an exponential decay. The end of an event is near if the positive slope decreased to a threshold. This threshold has to be as small as possible, but big enough that is sure that it will be triggered within the event. Figure 7b shows this on axis B.

These step trigger are not precise to define the start of an event. We chose the approach to linearize the step. Thus, the event start is the interception of the slope of the nearest inflection point to the nearest zero slope point before the step and the value of this zero slope point. Commonly, this interception point will be not on the signal, but represents a proper start time and start value.

### Event assigning

Equation (1) shows the thresholded distance matrix *C*_*i j*_ represents all meaningful combinations of events in the event sets *a* and *b*. By the threshold γ only combinations with a smaller distance survive in the matrix.

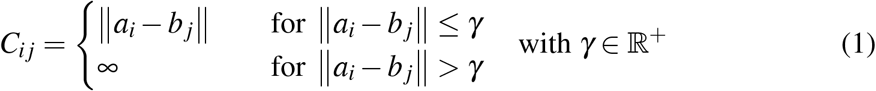

The assignment problem is given in equation (2), where *X* ∈ 𝔹^|*a*|×|*b*|^ is a sparse boolean matrix with *X*_*i j*_ = 1 if event *a*_*i*_ is assigned with event *b*_*j*_. The optimizer assigned all events as long as events in both sets are unassigned. This behavior can provoke fails. If an event that is not assignable will be assigned, could be one or more other events fail. By choosing a suitable maximum distance γ only near events will be assigned.

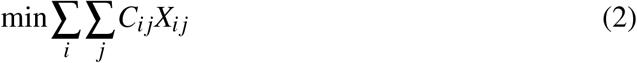

## Additional information

### Data and code availability statement

The artificial datasets generated during and analysed during the current study are available from the corresponding author on reasonable request.

## Acknowledgements

We thank Dr. Renato Frischknecht for providing the primary culture of mouse cortex cells. B. P. and A. F. were supported by grants (FE 464/12-1, FE 464/14-1) from the DFG (Deutsche Forschungsgesellschaft).

## Author contributions

Thomas Pircher and Bianca Pircher conceived of the presented ideas. Thomas Pircher wrote the code, performed the computations and technical experiments. Biological data and subsequent argumentation is contributed by Bianca Pircher and Andreas Feigenspan, who also supervised the findings of this work. Renato Frischknecht contributed to the experiments and provided the primary cortex cell culture. All authors discussed the results and contributed to the final manuscript.

## Code

The code of the described methods is available at:

- https://github.com/Digusil/eventsearch
- https://github.com/Digusil/openseapy

## References

1. Wall, E., Blaha, L. M., Paul, C. L., Cook, K. & Endert, A. Four Perspectives on Human Bias in Visual Analytics BT - Cognitive Biases in Visualizations. 29–42 (Springer International Publishing, Cham, 2018). URL https://doi.org/10.1007/978-3-319-95831-6{_}3.

2. Nickerson, R. S. Confirmation Bias: A Ubiquitous Phenomenon in Many Guises. Review of General Psychology 2, 175–220 (1998). URL https://doi.org/10.1037/1089-2680.2.2.175.

3. Cho, I. et al.. The Anchoring Effect in Decision-Making with Visual Analytics. In 2017 IEEE Conference on Visual Analytics Science and Technology (VAST), 116–126 (2017).

4. Chandola, V., Banerjee, A. & Kumar, V. Anomaly detection: A survey. ACM computing surveys (CSUR) 41, 1–58 (2009).

5. Faverjon, C. & Berezowski, J. Choosing the best algorithm for event detection based on the intended application: A conceptual framework for syndromic surveillance. Journal of biomedical informatics 85, 126–135 (2018).

6. Wheelwright, S., Makridakis, S. & Hyndman, R. J. Forecasting: methods and applications (John Wiley & Sons, 1998).

7. Box, G. E., Jenkins, G. M., Reinsel, G. C. & Ljung, G. M. Time series analysis: forecasting and control (John Wiley & Sons, 2015).

8. Kang, Y., Belušić, D. & Smith-Miles, K. Detecting and classifying events in noisy time series. Journal of the Atmospheric Sciences 71, 1090–1104 (2014).

9. Celisse, A., Marot, G., Pierre-Jean, M. & Rigaill, G. New efficient algorithms for multiple change-point detection with reproducing kernels. Computational Statistics & Data Analysis 128, 200–220 (2018).

10. McInnes, L., Healy, J., Saul, N. & Grossberger, L. Umap: Uniform manifold approximation and projection. The Journal of Open Source Software 3, 861 (2018).

11. Rothman, J. S. & Silver, R. A. Neuromatic: An integrated open-source software toolkit for acquisition, analysis and simulation of electrophysiological data. Frontiers in Neuroinformatics 12, 14 (2018). URL https://www.frontiersin.org/article/10.3389/fninf.2018.00014.

12. Kudoh, S. N. & Taguchi, T. A simple exploratory algorithm for the accurate and fast detection of spontaneous synaptic events. Biosensors and Bioelectronics 17, 773–782 (2002). URL https://www.sciencedirect.com/science/article/pii/S0956566302000532.

13. Muthmann, J.-O. et al.. Spike detection for large neural populations using high density multielectrode arrays. Frontiers in Neuroinformatics 9, 28 (2015). URL https://www.frontiersin.org/article/10.3389/fninf.2015.00028.

14. Frech, M. J., Pérez-León, J., Wässle, H. & Backus, K. H. Characterization of the spontaneous synaptic activity of amacrine cells in the mouse retina. Journal of neurophysiology 86, 1632–1643 (2001).

15. Nielsen, T. A., DiGregorio, D. A. & Silver, R. Modulation of glutamate mobility reveals the mechanism underlying slow-rising ampar epscs and the diffusion coefficient in the synaptic cleft. Neuron 42, 757–771 (2004). URL https://www.sciencedirect.com/science/article/pii/S0896627304002600.

16. Takekawa, T., Isomura, Y. & Fukai, T. Accurate spike sorting for multi-unit recordings. European Journal of Neuroscience 31, 263–272 (2010).

17. Feigenspan, A. & Babai, N. Functional properties of spontaneous excitatory currents and encoding of light/dark transitions in horizontal cells of the mouse retina. European Journal of Neuroscience 42, 2615–2632 (2015).

18. Babai, N. et al.. Signal transmission at invaginating cone photoreceptor synaptic contacts following deletion of the presynaptic cytomatrix protein bassoon in mouse retina. Acta physiologica (Oxford, England) e13241 (2018).

